# Adolescent social isolation creates a latent vulnerability in maternal care with intergenerational social consequences, rescued by experienced mothers

**DOI:** 10.64898/2026.04.14.718476

**Authors:** Jose Francis-Oliveira, Rinako Tanaka, Matthew Shen, Emily Cruvinel, Shin-ichi Kano, Minae Niwa

## Abstract

Early-life adversity (ELA) can produce lasting alterations on brain and behavior, yet its effects on maternal caregiving and intergenerational social outcomes remain poorly understood. Using a mouse model of mild adolescent social isolation, we demonstrated that females exposed to isolation in late adolescence exhibited progressive deficits in pup-directed maternal behaviors, including nursing, licking, nest-building, and pup retrieval, while self-directed behaviors remain intact. Offspring reared by these dams displayed deficits in adult social behavior, including reduced sociability, impaired social novelty recognition, and diminished social odor discrimination, without affecting non-social memory, anxiety-like behavior, or locomotor activity. Chemogenetic activation of the medial cingulate cortex (mCg) to prelimbic cortex (PrL) pathway in offspring of stressed dams rescued social deficits, whereas inhibition of this pathway in control mice recapitulated the impairments. Notably, co-housing stressed dams with experienced parous females during the early postpartum period restored pup-directed maternal behaviors and normalized offspring social outcomes, demonstrating that social experience can buffer the intergenerational consequences of ELA. Circuit analyses identified the mCg→PrL projections as a predominantly excitatory pathway critical for social behavior. Electrophysiological recordings revealed hypofunction of this circuit in offspring of stressed dams, which was ameliorated by postpartum co-housing with parous females. Together, these findings define a neural circuit linking adolescent psychosocial adversity to impaired maternal caregiving and intergenerational social dysfunction, and highlight social support from experienced mothers as a potent modulator of maternal and offspring behavioral outcomes.

**Significance Statement:** Adverse social experiences before motherhood can compromise maternal caregiving and transmit social vulnerability across generations, yet the underlying mechanisms remain poorly understood. Using a mouse model of subtle adolescent social isolation, we demonstrate that even mild isolation selectively impairs postpartum caregiving behaviors while sparing other behavioral domains, resulting in social deficits in adult offspring. We identify the midcingulate-prelimbic cortex circuit as a critical neural substrate mediating these effects and show that social support from experienced mothers during the early postpartum period restores both maternal behavior and offspring social outcomes. Together, these findings define a neural and social mechanism for the intergenerational effects of adolescent psychosocial adversity.

## Introduction

Maternal behavior is a fundamental social behavior that supports offspring survival and long-term health across species (1, 2). In mammals, early mother-infant attachment shapes both maternal mental health and offspring socioemotional development (3). Disruption of maternal care due to stress, psychiatric conditions, or limited social support can compromise offspring development (4–7). Early life adversity (ELA), including maltreatment, family dysfunction, trauma, and social deprivation, increases long-term risk for psychiatric and physical disorders and is associated with postpartum mood disturbances and impaired maternal care (8–13). However, the neural mechanisms by which ELA affects postpartum maternal care remain poorly understood, in part because most animal models rely on hormonal manipulations during pregnancy (14–18), which do not capture the lasting neurodevelopmental consequences of ELA on neural circuit function.

Adolescence is a sensitive period for brain and social development, marked by hormonal changes and refinement of circuits that regulate emotion and social behavior (19). Social isolation in rodents is a widely used model for psychosocial ELA (20–22), but most paradigms involve prolonged or severe isolation that produce immediate behavioral and endocrine deficits (23, 24). In contrast, we developed a model of mild adolescent social isolation that produces no detectable changes but generates a latent vulnerability that emerges during the postpartum period as impairments in maternal mood- and social-related behaviors (25, 26). This model enables investigation of how subclinical adolescent adversity programs maternal behavior and affects the next generation. Consistent with this framework, mild adolescent isolation is associated with prolonged dysregulation of the hypothalamic-pituitary-adrenal (HPA) axis and sustained corticosterone elevation specifically during the postpartum period, paralleling observations in women with postpartum depression following early adversity (26). Together, these findings define a developmental trajectory in which adolescent psychosocial adversity produces latent, context- dependent postpartum dysfunction, with stress-axis alterations serving as a downstream correlate rather that a sufficient explanatory mechanism.

Pregnancy induces lasting adaptations in the maternal brain, including structural remodeling of the anterior cingulate cortex (ACC) (27). In humans, the dorsal ACC (dACC), homologous to the rodent prelimbic cortex (PrL), is activated when healthy mothers respond to infant cues but shows reduced activation in mothers with caregiving difficulties (28, 29). The medial prefrontal cortex (mPFC), including the PrL, plays a central role in social cognition, emotional responses, and the expression of maternal behaviors (30–33). Within the rodent cingulate cortex, the anterior cingulate cortex [aCg; antero-posterior (AP) +0.0 mm] and mid-cingulate cortex (mCg; AP +1.0 mm) are structurally and functionally distinct, with the mCg implicated in attention, conflict monitoring, and social cognition (34–38). Although the role of the rodent mCg in maternal and social behaviors remains largely unexplored, its involvement in behavioral domains that are consistently disrupted following ELA suggests that it may represent a neural locus vulnerable to adversity-related dysfunction. Based on these observations, we hypothesize that disruption of cortico-cortical communication, particularly the mCg→PrL pathway, contributes to impaired maternal care and the intergenerational transmission of social deficits.

Maternal circuits remain plastic in adulthood, allowing social experience to shape caregiving behaviors. Mothers can learn maternal skills by observing experienced caregivers (39–44), highlighting the role of social learning in parenting. Social support may be particularly important for mothers with a history of ELA, who are at increased risk for postpartum mood disturbances that can compromise caregiving (45, 46). However, the mechanisms through which social support from experienced caregivers mitigates these deficits remain largely unknown.

In the present study, we show that mild adolescent social isolation creates a latent vulnerability in female mice that manifests during the postpartum period as impaired maternal care, mediated by reduced function of the mCg→PrL circuit. Dysfunction of this pathway drives social deficits in offspring, establishing a neural substrate for intergenerational transmission of ELA effects. Importantly, co-housing with parous females, defined here as females that have experienced at least one pregnancy and delivery, during the postpartum period ameliorates maternal behavioral deficits and rescues offspring social outcomes, revealing a mechanism through which social support can buffer the long-term consequences of early adversity.

## Results

### Adolescent social isolation impairs maternal caring behavior in the postpartum period

We first examined whether females exposed to social isolation during late adolescence exhibited alterations in maternal behavior after delivery. At postnatal day (P) 7, stressed dams spent significantly less time nursing (**Figure 1A**) and licking their pups (**Figure 1B**) compared to unstressed controls, resulting in a marked reduction in overall maternal caring behavior, defined as the combined duration of nursing and licking (**Figure 1C**). In contrast, the time spent climbing and digging (**Figure 1D**) as well as self-grooming (**Figure 1E**) did not differ between groups. Accordingly, total self-care behavior, defined as the combined duration of climbing/digging and self-grooming, was unchanged by adolescent isolation (**Figure 1F**). These findings indicate that adolescent social isolation selectively impairs maternal care toward offspring without broadly affecting self-directed maintenance behaviors. Consistent with this interpretation, nests built by stressed dams were smaller and less organized than those of unstressed controls (**Figure 1G**). Moreover, stressed dams required significantly more time to retrieve displaced pups to the nest (**Figure 1H**), further supporting impaired maternal responsiveness at one week postpartum.

**Figure 1.**
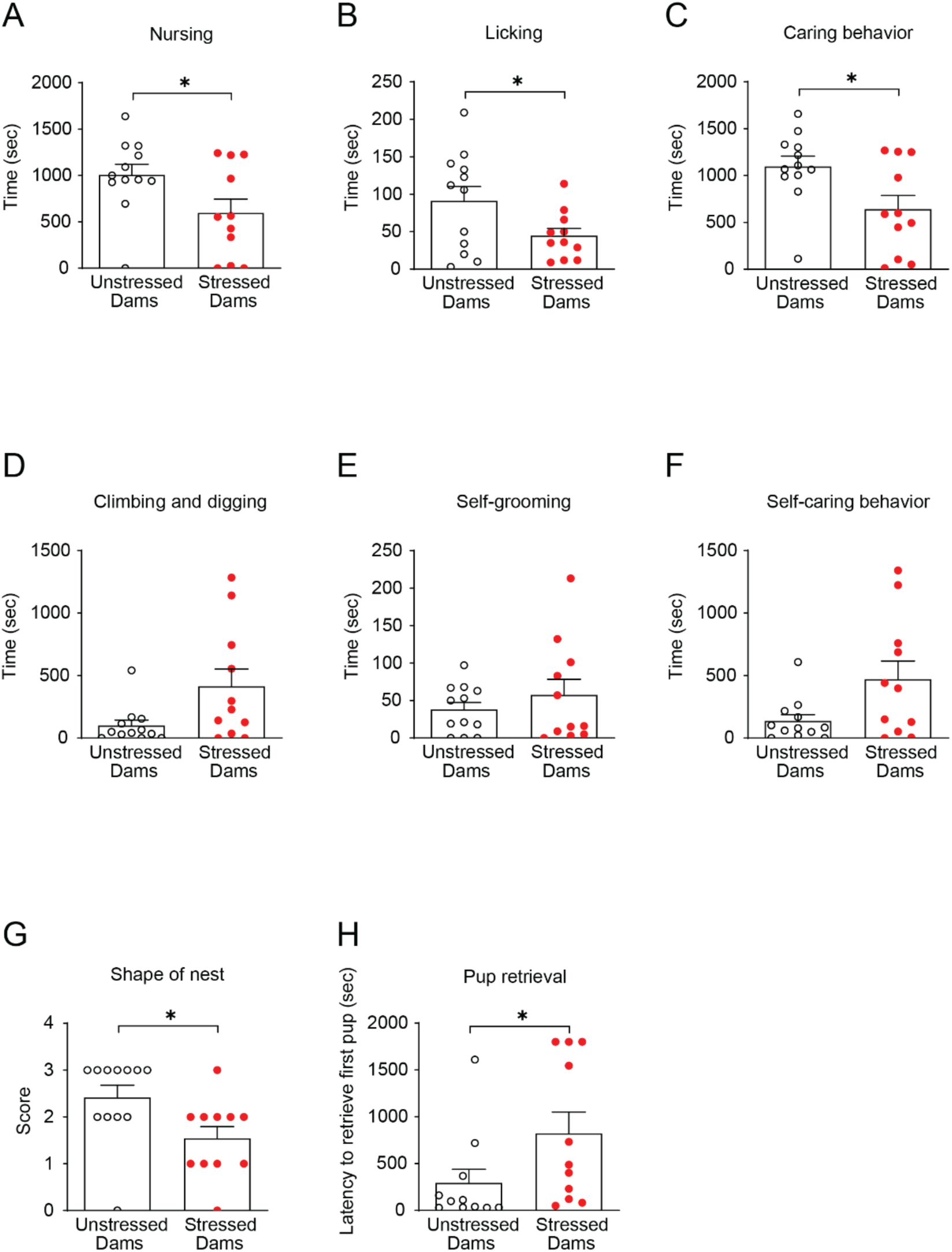
Adolescent social isolation impairs pup-directed maternal behavior without affecting self-directed behavior at 1-week postpartum. (A-B) Stressed dams showed reduced nursing (A) and pup-licking (B) behaviors. (C) Combined pup-directed care (nursing + licking) was significantly decreased in stressed dams. (D-E) Self-directed behaviors, including climbing/digging (D) and self-grooming (E), were unchanged. (F) Combined self-care behavior (climbing/digging + self-grooming) did not differ between groups. (G) Stressed dams built poorer-quality nests, reflected by lower nest scores. (H) Pup retrieval latency was significantly prolonged in stressed dams, indicating diminished maternal responsiveness. Data are shown as means ± SEM, N = 11-12 per group. *p < 0.05. Statistical tests: Student’s t-test (A, C), Welch’s t-test (B), and Mann-Whitney U (D-H) were applied as appropriate. Detailed statistical analyses are provided in **Supplementary Table 1**.

To determine whether these maternal deficits were evident immediately after birth, we assessed maternal behavior at P1. At this early stage, stressed dams showed only mild or partial impairments, limited to licking and pup retrieval behaviors (**Supplementary Figures 1A-H**). This pattern suggests that deficits in maternal care develop progressively over the first postpartum week, possibly reflecting delayed adaptation to the maternal care state.

Because maternal caregiving disruptions can influence offspring health, we evaluated the general well-being of the litters. Despite reduced maternal care, offspring reared by stressed dams (stressed offspring) exhibited body weight gain comparable to that of offspring reared by unstressed dams (unstressed offspring) (**Supplementary Figure 2A**). However, mortality rates were higher in litters reared by stressed dams, particularly during the early postnatal days (**Supplementary Figure 2B**). The total number of surviving pups per litter tended to be lower in stressed dams than in unstressed controls, and by P8, more stressed offspring had died compared with unstressed offspring; however, these differences were not statistically significant (**Supplementary Figure 2C**). These results indicate that adolescent social isolation leads to gradually emerging deficits in maternal caregiving during the first postpartum week, which contribute to increased early pup mortality despite apparently normal growth among surviving offspring.

### Adolescent isolation-induced maternal deficits lead to impaired social behavior in adult offspring

We next examined whether postpartum maternal caregiving deficits induced by adolescent social isolation affected behavioral outcomes in offspring during adulthood (>P56). In the social interaction test (SIT), stressed offspring exhibited reduced sociability and impaired social novelty recognition compared with unstressed controls (**Figures 2A and 2B**). In the social olfactory recognition test (SORT), stressed offspring showed markedly reduced investigation toward both familiar social cues from the test mouse’s home cage and novel social cues from the stranger’s cage (**Figure 2C**), indicating broad impairments in social cue recognition. In contrast, stressed offspring showed no differences from unstressed offspring in the novel object recognition test (NORT), open field test (OFT), elevated plus maze test (EPM), or light-dark box test (LDB) (**Figures 2D-G**). Locomotor activity in the OFT and discrimination index during the NORT training session were also comparable between groups (**Supplementary Figures 3A and 3B**). Consistent with heightened stress sensitivity, serum corticosterone levels were significantly elevated in adult stressed offspring relative to unstressed controls (**Supplementary Figure 4**).

**Figure 2.**
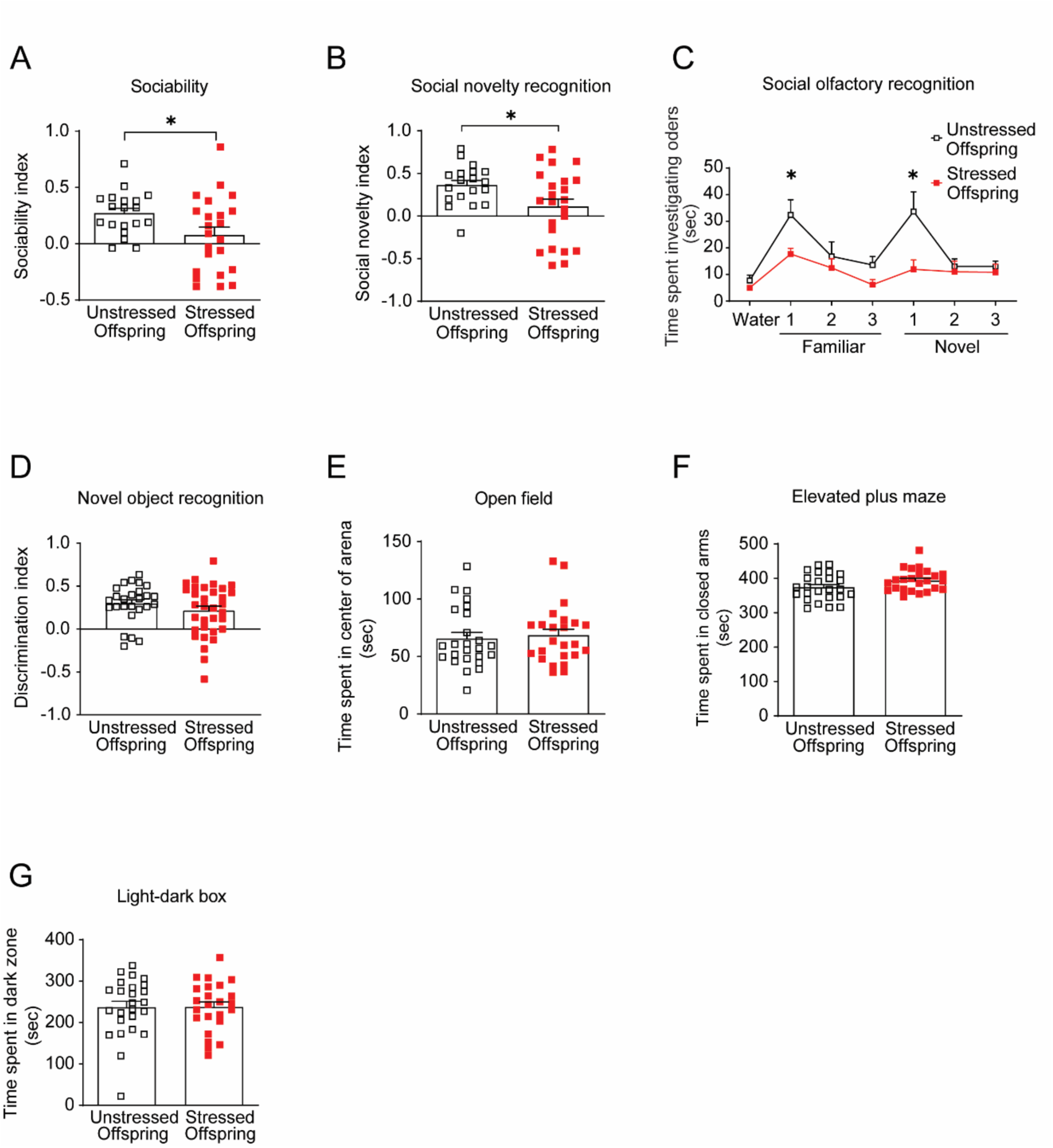
Offspring reared by stressed dams show selective social behavior deficits. (A) In the three-chamber social interaction test (SIT), stressed offspring showed reduced sociability compared with unstressed offspring. (B) Stressed offspring also displayed impaired social novelty recognition in the SIT. (C) In the social olfactory recognition test (SORT), stressed offspring exhibited diminished investigation toward both familiar and novel social odors. (D) Performance in the novel object recognition test (NORT) was unaffected by maternal adversity. (E) In the open field test (ORT), time spent in the center did not differ between groups. (F and G) Anxiety-related behaviors in the elevated plus maze (EPM) and light-dark box (LDB) showed no significant differences between groups. Together, these results indicate that impaired maternal behavior in stressed dams selectively disrupts offspring social behavior while sparing cognitive and anxiety-related domains. Data are shown as means ± SEM, N = 18-32 per group. *p < 0.05. Statistical tests: Welch’s t-test (A, B), two-way mixed ANOVA (C), Mann-Whitney U (D, G), and Student’s t-test (E, F), and were applied as appropriate. Detailed statistics are provided in **Supplementary Table 1**.

These findings suggest that adolescent isolation-induced deficits in maternal caregiving selectively impair social behavior in adult offspring without affecting general locomotion, anxiety-related behavior, or non-social memory. The persistent elevation in corticosterone levels indicates long-lasting dysregulation of HPA axis function that may contribute to the observed social deficits. Collectively, these results support the idea that disrupted maternal care during the early postnatal period can produce long-lasting, domain-specific alterations in offspring social behavior and stress physiology.

### The mCg→PrL pathway regulates social behavior and mediates the effects of poor maternal care

Given the established role of the PrL in regulating social behavior (28) and our hypothesis that cortico-cortical connectivity serves as a key circuit substrate for social behavior, we next investigated whether the mCg→PrL pathway plays a critical role in social behaviors in offspring.

To manipulate this pathway, we employed Designer Receptors Exclusively Activated by Designer Drugs (DREADDs), expressing engineered G-protein coupled receptors [M3 muscarinic receptor (hM3Dq; excitatory) and hM4Di (inhibitory)] selectively activated by the synthetic ligand clozapine-N-oxide (CNO) (47).

Excitation of the mCg→PrL pathway in stressed offspring restored sociability and social novelty recognition in the SIT (**Figures 3A-C**). Conversely, inhibition of the pathway in unstressed offspring induced behavioral deficits similar to those observed in stressed offspring (**Figures 3D-F**). These results indicate that hypofunction of the mCg→PrL pathway contributes to social behavioral deficits in stressed offspring and that the pathway is functionally required for normal social behavior.

**Figure 3.**
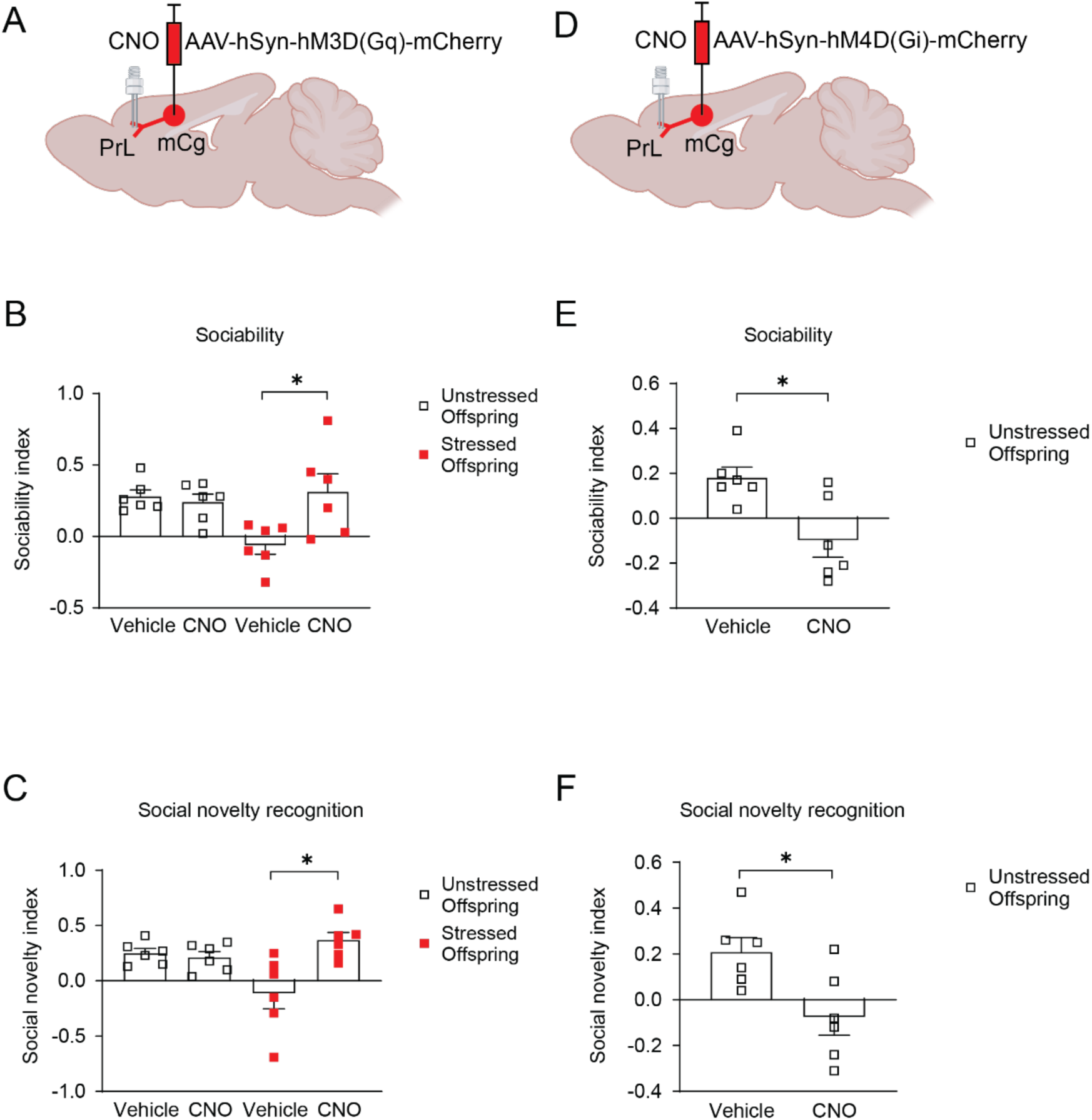
Chemogenetic modulation of mCg→PrL pathway rescues or induces social deficits. (A) Schematic illustrating virus delivery of AAV-hSyn-hM3D(Gq)-mCherry into the mCg and placement of guide cannulas into the PrL. (B and C) Activation of M3Dq-expressiong mCg→PrL neurons via CNO in stressed offspring rescued sociability and social novelty recognition deficits. (D) Schematic illustrating virus delivery of AAV-hSyn-hM4D(Gi)-mCherry into the mCg and placement of guide cannulas into the PrL. (E and F) Inhibition of the same circuit in unstressed offspring using M4Di receptors induced social deficits. Data are shown as means ± SEM, N = 6 per group. *p < 0.05. Statistical tests: Two-way ANOVA (B-C) and Student’s t-test (E-F) were applied. Detailed statistics are provided in **Supplementary Table 1**.

### Co-housing with parous females rescues pup-directed maternal behaviors and offspring social deficits

Parous mice exhibited significantly higher levels of maternal care than virgin mice, even when caring for foster pups (**Supplemental Figures 5A-G**). Specifically, parous females spent more time nursing (**Supplemental Figure 5A**) and licking pups (**Supplemental Figure 5B**), resulting in greater overall caring behavior, defined as the combined duration of nursing and licking (**Supplemental Figure 5C**). In contrast, self-directed behaviors, including climbing and digging (**Supplemental Figure 5D**) and self-grooming (**Supplemental Figure 5E**), did not differ between groups. Accordingly, total self-care behavior, defined as the combined duration of climbing/digging and self-grooming, was comparable between parous and virgin females (**Supplemental Figure 5F**). Pup retrieval behavior was significantly increased in parous females relative to virgins (**Supplemental Figure 5G**). Together, these findings indicate that pregnancy and delivery selectively enhance offspring-directed maternal behaviors without broadly affecting self-directed maintenance activities.

Based on these observations, we next investigated whether social exposure to experienced mothers could rescue maternal deficits. Stressed dams were co-housed with parous females from P0 to P7, the day following completion of maternal behavioral testing. During co-housing, pups remained with the stressed dams and were accessible to both stressed and parous females, allowing observation and potential social learning. Immediately prior to maternal behavioral testing, the parous females were temporarily removed so that measured behaviors reflected the stressed dams’ responses to their own pups rather that direct interactions with the parous females. This experimental design is summarized in **Figure 4A**. Co-housing from P0 to P7 improved nest-building and rescued deficits in pup-directed caring and retrieval behaviors in stressed dams, while self-directed behaviors remained unchanged (**Figures 4B-I**). In contrast, co-housing had no effect on maternal behavior in unstressed dams (**Figures 4B-I**).

**Figure 4.**
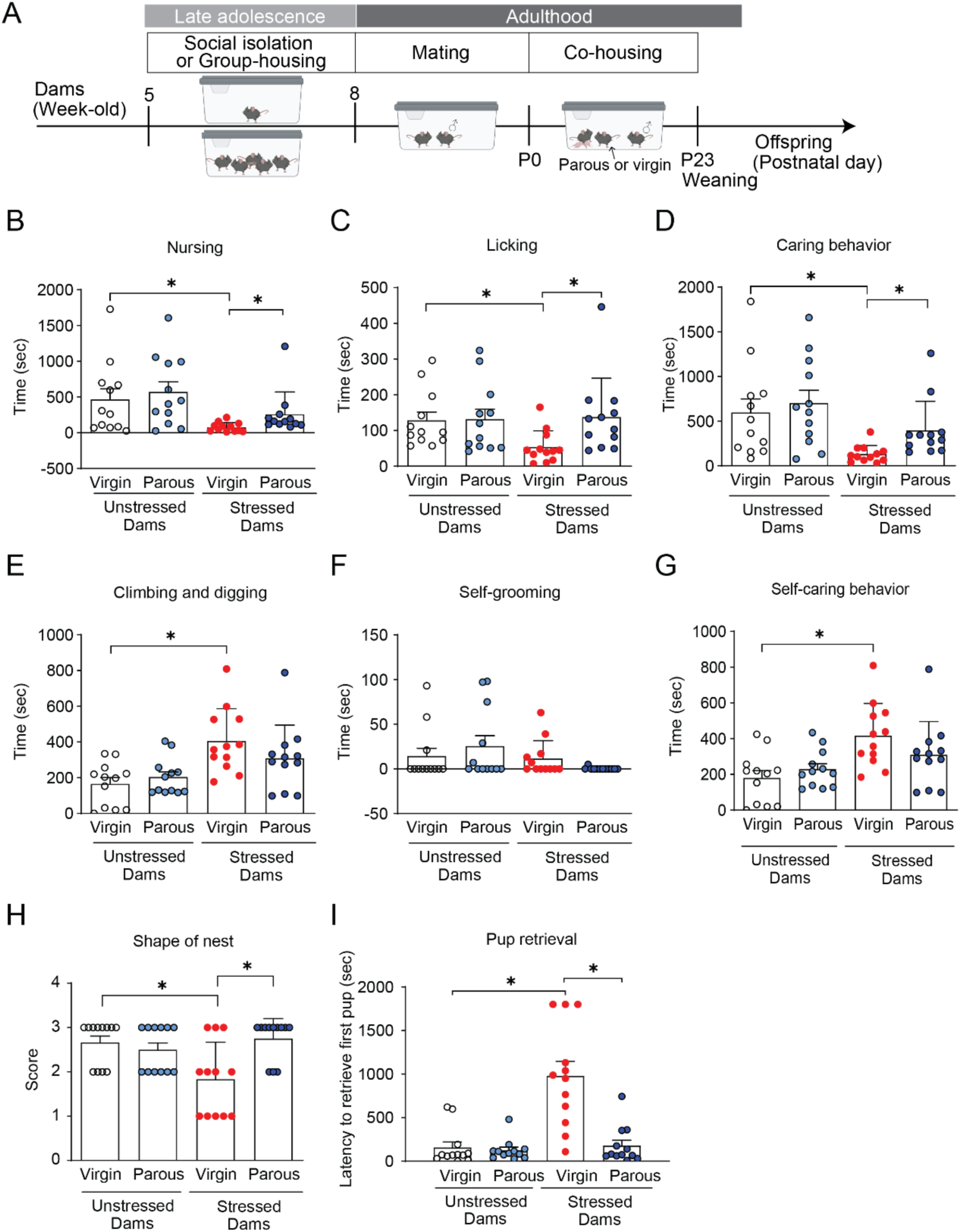
Co-housing with parous females rescues impaired maternal caregiving in stressed dams. (A) Experimental design of co-housing with parous females. (B-D) Pup-directed behaviors were increased in stressed dams co-housed with parous, but not virgin, females. (E-G) Self-care behaviors were not affected by co-housing conditions. (H and I) Nest quality and pup retrieval performance were improved following co-housing with parous females. Data are shown as means ± SEM, N = 12 per group. *p < 0.05. Statistical tests: All non-normal data were analyzed using pairwise planned comparisons in Mann-Whitney U tests. Detailed statistics are provided in **Supplementary Table 1**.

Importantly, this maternal behavioral rescue translated into improved offspring behavioral outcomes. Stressed offspring reared by dams co-housed with parous females during the postpartum period exhibited restored sociability and social novelty recognition in the SIT (**Figures 5A and 5B**). These findings indicate that social support from experienced mothers during the early postpartum period selectively mitigates pup-directed maternal deficits and ameliorates intergenerational social behavioral impairments.

**Figure 5.**
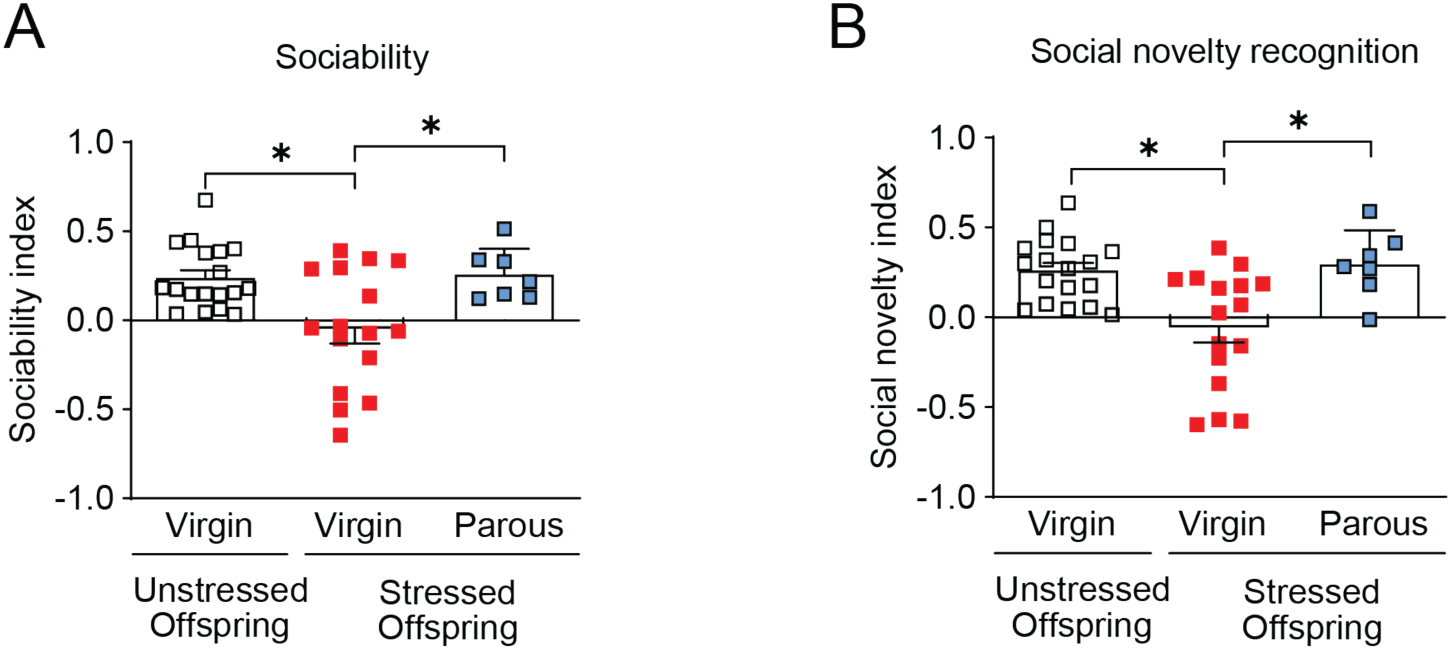
Co-housing with parous mice rescues social deficits in stressed offspring. (A) Sociability index was decreased in stressed offspring but rescued by co-housing with parous females from birth to weaning. (B) Social novelty recognition followed a similar pattern and was rescued by parous co-housing. Data are shown as means ± SEM. Sample size for sociability index is N = 7-18 per group. *p < 0.05. Statistical tests: Kruskal-Wallis test (A) and Welch’s one-way ANOVA (B) were applied. Detailed statistics are provided in **Supplementary Table 1**.

### Excitatory mCg→PrL firing is reduced in stressed offspring and restored by co-housing with parous females

Building on evidence of cortico-cortical connectivity between the mCg and PrL in mice, we performed retrograde viral tracing to identify upstream cortical inputs to the PrL. Consistent with our prior identification of mCg→PrL projections (36), we focused on further characterizing this pathway. Following retrograde viral injection into the PrL, the majority of labeled mCg cells co-expressed CaMKII, a marker of excitatory neurons, whereas only a small fraction co-localized with GABA, an inhibitory marker (**Figures 6A-C**). Furthermore, injection of a CaMKII-promoter-driven retrograde virus into the PrL labeled numerous mCg cells (**Figure 6D**), confirming that the mCg→PrL pathway is predominantly excitatory.

**Figure 6.**
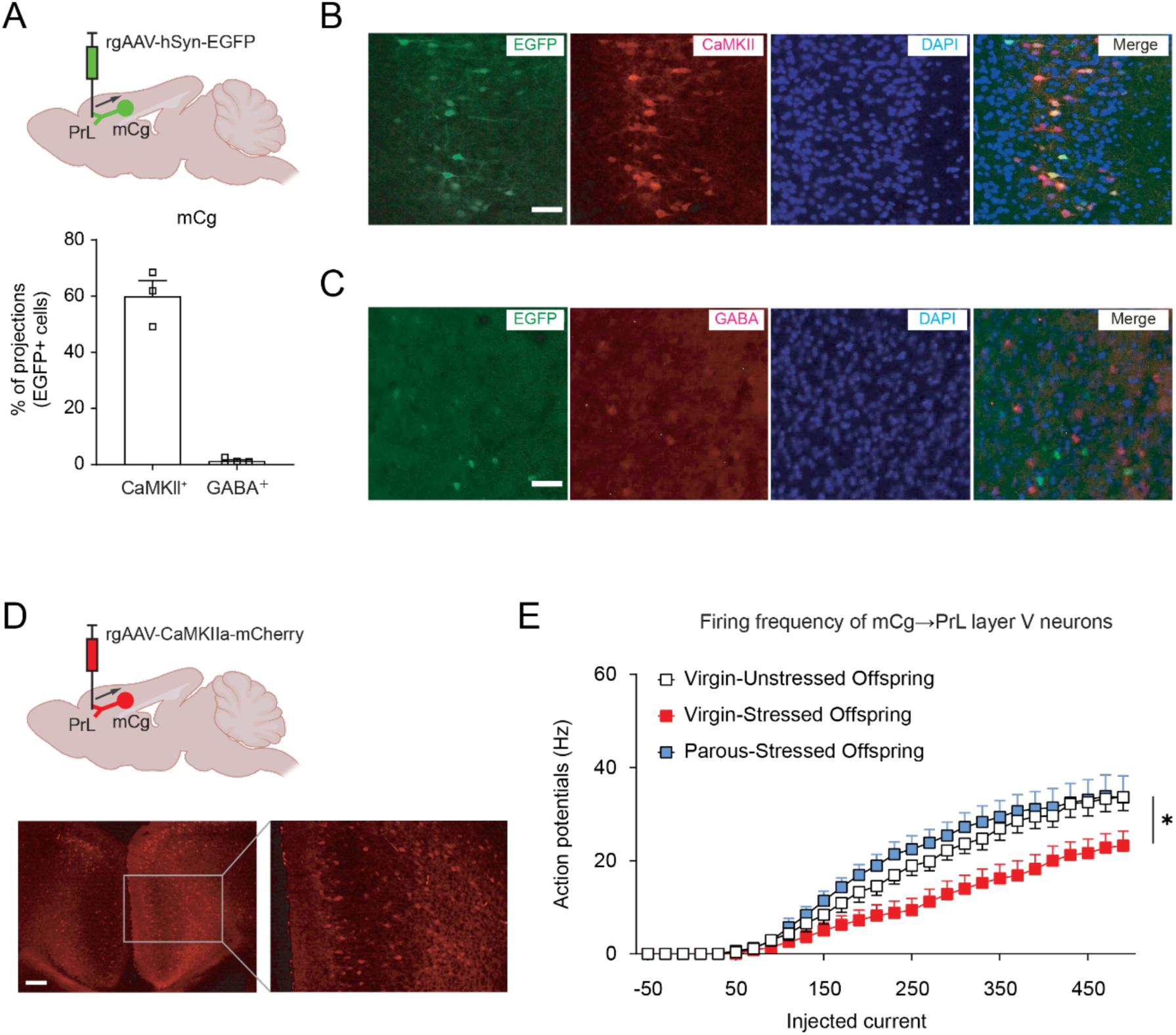
The mCg→PrL pathway is predominantly excitatory and shows reduced firing in stressed offspring that is restored by co-housing. (A) Left: Schematic illustrating virus delivery of rgAAV-hSyn-EGFP into the PrL. Right: Quantification of EGFP-labeled mCg neurons retrogradely labeled from PrL co-localized with CaMKII or GABA. Data are shown as means ± SEM, N = 3 per group. Each dot represents an average of three slices per animal. (B and C) Representative immunohistochemistry images of EGFP-labeled mCg neurons retrogradely labeled from PrL co-localized with CaMKII or GABA. Scale bars: 50 µm. (D) Schematic image and representative mCg image showing mCherry-labeled neurons following injection of a CaMKll promoter-driven retrograde virus into the PrL, confirming the excitatory nature of the mCg→PrL circuit. Scale bars: 100 µm. (E) Firing frequency of action potentials in layer V mCg→PrL neurons was reduced in stressed offspring, and rescued in stressed offspring co-housed with parous mice. Sample size for firing frequency is 10-13 cells per group, obtained from 4 different mice in each group. *p < 0.05. Statistical tests: Two-way mixed ANOVA (C) was used. Detailed statistics are provided in Supplementary Table 1.

We next assessed the electrophysiological properties of mCg→PrL neurons in adult offspring. Because cortico-cortical projection neurons from the mCg to the PrL are predominantly located in layer V (48, 49), we focused our recordings on neurons in this layer. Whole-cell patch-clamp recordings revealed a significant reduction in firing frequency in stressed offspring compared with unstressed controls (**Figure 6E**), indicating hypofunction of the mCg→PrL pathway. In contrast, rheobase and resting membrane potential were unchanged (**Supplemental Figures 6A and 6B**), suggesting that intrinsic membrane excitability was not altered. Analysis of spontaneous excitatory and inhibitory postsynaptic currents showed no significant differences in amplitude or frequency between groups (**Supplemental Figures 6C-F**), indicating that presynaptic input to mCg→PrL neurons remains intact. Collectively, these data suggest that impaired maternal care reduces the output activity of mCg→PrL neurons without affecting their intrinsic excitability or upstream synaptic inputs. Notably, co-housing with parous females restored firing activity in these neurons (**Figure 6E**), corresponding with normalized social behaviors in adult offspring (**Figure 5**).

## Discussion

Social isolation during adolescence establishes a latent vulnerability in female mice, selectively impairing postpartum pup-directed behaviors, including licking, nursing, nest-building, and pup retrieval. These specific maternal deficit traits are transmitted to the next generation, causing persistent social deficits in adult offspring, while sparing other behavioral domains. Our research revealed that the mCg→PrL projection is a critical excitatory pathway underlying these intergenerational effects. Crucially, social contact with experienced parous mothers during the early postpartum period selectively restores maternal care and rescues offspring’s social behavior. This demonstrates a causal link between maternal experience, circuit function, and offspring development. These findings underscore the dual importance of adolescent psychosocial experiences and the postpartum social environment in shaping maternal behavior and offspring social trajectories.

The prefrontal cortex constitutes a core substrate for maternal and social behavior. Our data indicate that adolescent social isolation reduces the firing output of an excitatory cortico-cortical pathway in the prefrontal cortex (mCg→PrL), while presynaptic inputs remain largely intact. This is consistent with prior reports that ELA selectively remodels excitatory cortico-cortical circuits, with behavioral specificity reflecting vulnerability of defined projections (25, 32, 50, 51). Candidate mechanisms include altered ion channel expression and modulation of conductance-regulating signaling pathways, reminiscent of TRPM3-dependent regulation in social hierarchy circuits (50). Functional manipulation sufficiently rescued social deficits by restoring activity within this pathway, while its inhibition was confirmed to mimic adversity-induced impairments, emphasizing a causal role for excitatory cortico-cortical dynamics in integrating maternal cues and guiding offspring social outcomes.

The selectivity of behavioral changes in offspring highlights the critical influence of early maternal care on social development (52, 53). Offspring exposed to suboptimal maternal care exhibit elevated corticosterone levels while maintaining normal locomotion, anxiety responses, and non-social memory, suggesting persistent stress axis dysregulation (6, 54). This pattern aligns with prior evidence linking deficits in tactile and nursing stimulation to impaired HPA axis negative feedback and abnormal stress responsivity (26, 55, 56). While oxytocinergic signaling may also contribute to social vulnerability, its precise regulation by maternal care remains unexplored (57–60). Our findings suggest that maternal deficits shape the development of social circuits without broadly impairing other behavioral domains, providing a framework for understanding how ELA confers selective vulnerability to social dysfunction across generations.

Communal rearing with experienced parous dams selectively improves maternal behavior in stressed females, demonstrating the plasticity of maternal circuits (61, 62). Mechanistically, this effect may arise through three complementary pathways: (1) Social support – affiliative presence reduces psychological stress and enhances caregiving motivation (63); (2) Behavioral modeling – stressed dams learn maternal behaviors by observing parous partners, leading to sustained improvements even when the model is absent (40); (3) Stress buffering – enhancing caregiving capacity through normalization of corticosterone levels and restoration of mCg→PrL neuronal activity (64, 65). These results align with prior work demonstrating that maternal behavior can be socially transmitted in mice and other mammals (39, 40, 66), emphasizing the translational potential of supportive social environments to buffer adversity.

Collectively, our study elucidated a mechanistic framework linking adolescent psychosocial adversity to postpartum maternal dysfunction and selective intergenerational social deficits. By identifying the mCg→PrL pathway as a critical substrate for experience-dependent regulation of social behavior, these findings provide a basis for interventions to mitigate the persistent effects of ELA. Future studies should elucidate the molecular and synaptic mechanisms underlying circuit hypofunction, investigate the potential for transgenerational epigenetic modifications, and explore the efficacy of structured social support interventions across diverse mammalian models. Understanding these processes offers insight into the pathogenesis of social dysfunction in humans and contributes to developing strategies that promote resilience to ELA.

## Materials and Methods

### Mice

C57BL/6J (B6J; JAX 000664) and B6;129S-Slc17a7tm1.1(cre)Hze/J (Vglut1-Cre; 023527) mice were obtained from the Jackson Laboratory. All mice were backcrossed for more than ten generations onto the C57BL/6J genetic background. Mice were maintained under controlled environmental conditions (23 ± 3 °C; 40 ± 5% humidity) with a 12-hour light/dark cycle (lights on at 6 a.m., lights off at 6 p.m.). Animals were housed in wire-topped plastic cages (20 × 31 × 13 cm) with ad libitum access to food and water. All experimental procedures adhered to the National Institutes of Health Guidelines for the Care and Use of Laboratory Animals and were approved by the Institutional Animal Care and Use Committee at the University of Alabama at Birmingham (IACUC-21547). Both sexes were included to ensure generalizability, but sex differences were not analyzed because the study was primarily designed to test maternal and offspring outcomes and was not powered for sex-specific comparisons.

#### Unstressed and Stressed Dams; Unstressed and Stressed Offspring

Virgin female mice were either group-housed or socially isolated from five to eight weeks of age to induce psychosocial adversity. At eight weeks of age, all females were mated with healthy B6J males. Pregnancy was monitored daily, and the day of delivery was designated as P0. These mice were designated as stressed dams. Age-matched controls (unstressed dams) were group-housed (≥3 per cage) in wire-topped plastic cages during the same period. Pups raised by unstressed dams were designated unstressed offspring, and pups raised by stressed dams were designated stressed offspring. All offspring were weaned at P23 and group-housed thereafter until adulthood (>P56), when experimental procedures were performed. Both sexes were used in all offspring experiments.

#### Parous Mice

Female nulliparous mice were group-housed and mated, and following weaning of their litters, they remained group-housed until use. These dams, with only one prior delivery and maternal experience, are referred to as “parous females.” To assess the effects of co-housing on maternal behavior, either a virgin or parous female was introduced into the cages of unstressed or stressed dams with their pups immediately after delivery (P0) (**Fig. 4A**). Co-housed females remained together for the first postpartum week, during which maternal behaviors were evaluated in both the co-housed females and the experimental dams. To examine the long-lasting impact of co-housing with parous females on offspring social behavior, virgin or parous females were co-housed with unstressed or stressed dams and their pups from birth until weaning (P23). The introduced females were unfamiliar with the focal dams prior to co-housing. All co-housed females were age-matched and had comparable social histories, except for prior pregnancy and delivery. For parous females, at least three to five weeks elapsed between weaning of their previous litter and introduction to the focal dams, ensuring recovery and stabilization of maternal experience.

### Behavioral testing overview

Basal maternal behavior and pup retrieval tests were conducted on P1 and P7, with both tests performed at each time point. Unstressed and stressed dams co-housed with either parous or virgin females were evaluated on P7. Separate cohorts of dams were used for P1 and P7, and data from each timepoint were analyzed independently.

Behavioral assessments for adult offspring were conducted according to the experimental timeline. The SIT included a habituation phase on P53-55 and the test phase on P56, followed by the open field test on P57, the elevated plus maze test on P58, the light-dark box test on P59, and brain/blood sampling on P60. Separate cohorts of mice were used for the novel object recognition test on P56 and the social olfactory recognition test on P56.

Behavioral data were analyzed using EthoVision XT (versions 15 and 17; Noldus Information Technology) for automated tracking, and manual measurements were used for the basal maternal behavior and pup retrieval tests.

#### Basal Maternal Behavior Test

The basal maternal behavior test was conducted as previously described with minor modifications (67, 68). Basal maternal behaviors were assessed in the home cage to evaluate spontaneous caregiving activity in dams under undisturbed conditions. Each dam was observed for 30 min, and all behaviors were recorded for subsequent manual scoring by an observer blinded to experimental conditions. Nest quality was scored on a 4-point scale (0–3) based on structural completeness and pup coverage, as follows: 0 = no nest; scattered bedding with no structure; 1 = shallow or incomplete nest structure; 2 = well-defined nest with partial enclosure of pups; 3 = fully enclosed nest with pups gathered and covered. The duration (in seconds) of pup-directed and non-pup-directed behaviors was quantified as follows: Pup-directed behaviors: *Licking:* the dam licks or grooms any part of the pup’s body, primarily the anogenital region. *Nursing:* the dam displays a crouching or arched-back posture with legs extended to cover the litter. *Caring behavior:* total duration of licking and nursing combined. Non-pup-directed behaviors: *Climbing and digging:* pushing or displacing bedding material with the snout or forepaws. *Self-grooming:* licking or scratching any part of the dam’s own body. *Self-care:* total duration of climbing/digging and self-grooming combined. All behaviors were manually scored from video recordings by an experimenter blind to group assignments.

#### Pup Retrieval Test

Maternal responsiveness was evaluated using the pup retrieval test, as previously described with minor modifications (40, 59, 69). Pups were temporarily removed from the home cage, and dams were allowed to habituate for 30 minutes prior to testing. Three pups from the dam’s own litter were placed individually in separate corners of the home cage, equidistant from the nest site. In the P7 condition, to prevent spontaneous pup movement, transparent cups were placed in three corners of the cage, and one pup from the dam’s own litter was placed in each cup. The latency to retrieve each pup and return it to the nest was recorded. The dam was given up to 30 minutes to complete the retrieval of all three pups. The primary measure was the latency to retrieve the first pup (in seconds). If the dam failed to retrieve any pup within 30 min, a latency of 1800 s was assigned for statistical analysis. All trials were video recorded and scored by an observer blind to the experimental conditions.

#### Social Interaction Test (SIT)

The SIT was performed in adult offspring as previously described with minor modifications (25, 26, 70–72). The test consisted of two sequential trials: the Sociability Trial (S-trial) and the Social Novelty Trial (SN-trial), which were designed to assess progressively complex social behaviors. The testing apparatus consisted of a three-chamber box (60 cm × 40 cm × 35 cm) divided into left, middle, and right chambers, with small, barred cages positioned in the left and right compartments. The subject mouse was allowed to habituate to the three-chamber environment for 10 minutes daily over three consecutive days before testing. On the test day, each mouse was placed in the middle chamber and allowed to habituate for 10 minutes before being confined to the middle chamber. The doors were then closed to create three separate chambers, and the subject mouse returned to the middle chamber for 5 minutes. During this period, a novel mouse was introduced into one of the barred cages, while a mouse-shaped object was placed in the other barred cage. In the S-trial, the subject mouse encountered the mouse in the barred cage and the object in the other barred cage for 10 minutes. At the 10-minute mark, the test mice were again restricted from entering the right and left chambers for 5 minutes, and the mouse-shaped object was replaced with another novel mouse. During the SN-trial, the subject mouse encountered a familiar mouse in one barred cage and a novel mouse in the other barred cage for 10 minutes. Familiar and novel mice were chosen to be age- and sex-matched healthy mice. Behavioral data were recorded and analyzed using Ethovision XT 17 software (Noldus). Nose contact and proximity within a 1cm boundary zone surrounding each cage were automatically detected, and the time spent sniffing each cage were quantified. Sociability and social novelty indexes were calculated as follows: (time interacting with mouse or novel mouse cage – time interacting with an object or familiar mouse cage) / (time interacting with a mouse or novel mouse cage + time interacting with an object or familiar mouse cage). An index of 0 indicated no preference. Positive indexes indicate increased sociability or social novelty behavior, whereas negative indexes indicated social avoidance or deficits in social novelty behavior.

#### Social Olfactory Recognition Test (SORT)

The SORT was conducted in adult offspring to assess social odor recognition and discrimination, as previously described with minor modifications (73). Each mouse was habituated to a clean testing cage, identical in dimensions to its home cage, for 30 min before testing. Odor stimuli were presented using cotton balls soaked with odor solutions and placed in open 15 mL conical tubes secured at one corner of the cage using a custom plastic holder. Three odor conditions were used: (1) MilliQ water, (2) familiar scent, generated by agitating soiled bedding from the subject’s own home cage in MilliQ water, and (3) stranger scent, prepared from soiled bedding collected from an age- and sex-matched unfamiliar mouse housed in a separate cage with no prior contact. Each odor type was presented in three consecutive 2-min trials with no inter-trial intervals. Cotton balls were replaced between trials to prevent cross-contamination and ensure consistent odor presentation. The testing sequence was water → three sessions of familiar scent → three sessions of stranger scent. Olfactory investigation was defined as direct nasal contact or proximity (≤ 2.54 cm) to the cotton ball. Investigation time was recorded for each trial and quantified as total exploration duration by an observer blinded to the experimental group. Animals displaying little exploratory behavior (<10 s cumulative investigation across all trials) were excluded from subsequent analysis.

#### Novel Object Recognition Test (NORT)

The NORT was conducted in adult offspring as previously described with minor modifications (74, 75). Each trial consisted of three phases: habituation, training, and testing, each lasting 10 minutes. During the habituation phase on postnatal days 52-54, the subject mouse was introduced into a chamber apparatus (40 cm × 40 cm × 42 cm) and allowed to habituate for 10 minutes per day over the course of three days. On postnatal day 55, during the training phase, the mouse was once again habituated for 10 minutes in the chamber apparatus. Following this, the mouse was briefly removed and then reintroduced into the chamber, which contained two identical cone objects, for 2 minutes. Data during the training phase were collected over the subsequent 10 minutes. On postnatal day 56, the test phase began with a 10-minute habituation in the chamber apparatus. Subsequently, the mouse was briefly removed from the chamber.

During this brief removal, one of the two familiar cone objects was replaced with a novel pyramid object. The mouse was then returned to the chamber for 2 minutes, after which data collection took place over the following 10 minutes. All data were manually analyzed by reviewing video recordings to identify instances of nose contact or physical interactions within a 1cm boundary zone around the objects. Stopwatch timers were used to measure the time spent sniffing. The discrimination index was calculated as follows: (time sniffing the novel object – time sniffing the familiar object / (time sniffing the novel object + time sniffing the familiar object). An index of 0 indicated no recognition of the novel object, positive values indicated increased recognition of the novel object, and negative values indicated deficits in recognizing the novel object.

#### Open Field Test (OFT)

The OFT was conducted in adult offspring to assess general locomotor activity, as previously described with minor modifications (76). Each mouse was placed in the center of a square open-field arena (40 cm × 40 cm × 42 cm) constructed from opaque material to prevent external visual distractions. The mice were allowed to explore the arena freely for 10 min under standardized room lighting and ambient noise conditions. The total distance traveled was automatically recorded and analyzed using EthoVision XT software.

#### Elevated Plus Maze Test (EPM)

Anxiety-like behavior was evaluated in adult offspring using the EPM, as previously described with minor modifications (77–79). The maze consisted of two open arms (40 × 10 cm) and two closed arms (40 × 10 cm, enclosed by 20-cm-high opaque gray walls) elevated 50 cm above the floor. The apparatus was positioned in a quiet, dimly lit room to minimize environmental stressors. Each mouse was placed in the central platform facing an open arm and allowed to explore the maze freely for 10 min. Behavioral parameters measured included time spent in open versus closed arms, number of entries into each arm, and total distance traveled. All data were automatically recorded and analyzed using EthoVision XT software. Time spent in closed arms was used as the primary index of anxiety-like behavior. The experimenter was positioned outside the line of sight of the animal to avoid introducing stress-related artifacts.

#### Light-Dark Box test (LDB)

The LDB was conducted in adult offspring to assess anxiety-like behavior, as previously described with minor modifications (80). The apparatus consisted of two compartments: a dark chamber (opaque black walls and lid, impervious to visible light) and a light chamber (opaque walls, illuminated under standard room lighting). The chambers were connected by a small opening, allowing free movement between compartments. Each mouse was placed in the center of either the light or dark compartment (facing away from the opening) and allowed to explore for 10 min. Behavioral measures included total time spent in each compartment, number of transitions between compartments, and latency to first enter the dark chamber. Time spent in the dark chamber was considered a measure of anxiety-like behavior, with greater time indicating increased anxiety. All behavior was automatically tracked and analyzed using EthoVision XT software. Lighting and ambient noise were controlled to minimize environmental variability.

#### DREADD manipulation of the mCg→PrL pathway

To selectively manipulate the mCg→PrL pathway, we employed DREADDs, which are engineered G-protein coupled receptors activated exclusively by synthetic ligands such as CNO (47). At P35, mice were anesthetized with isoflurane (5% induction, 1 - 2.5% maintenance) and placed in a stereotaxic apparatus. Bilateral stereotaxic injections of 400 nL AAV-hSyn-hM3D(Gq)-mCherry (4.3 × 10^12^ genomic copies/mL; #50474, Addgene) or AAV-hSyn-hM4D(Gi)-mCherry (4.0 × 10^12^ genomic copies/mL; #50475, Addgene) were delivered into the mCg [coordinates from bregma: AP +0.2 mm, medial-lateral (ML) ±0.3 mm, dorsal-ventral (DV) −1.7 mm] at a rate of 0.1 μL/min. The injection cannula was left in place for 5 minutes post-injection to allow diffusion and prevent backflow. hM3Dq and hM4Di were used to induce neuronal excitation and inhibition, respectively (81). During the same surgery, bilateral guide cannulas (C235G-1.0/SPC, 26-gauge, Protech International) were implanted targeting the PrL (AP +2.34 mm, DV −1.45 mm, 0.5 mm above the intended microinjection site at DV −1.95 mm). Dummy cannulas were inserted to maintain guide patency until P56. At P56, internal cannula extending 0.5 mm beyond the guides were used for microinjections. Neuronal activity was manipulated 30 minutes prior to SIT via bilateral microinjection of CNO (300 μM, 300 nL) or vehicle [1% dimethyl sulfoxide (DMSO) in artificial cerebrospinal fluid (aCSF)] into the PrL (AP +2.34 mm, ML ±0.4 mm, DV −1.95 mm) using microsyringe pumps. Viral expression in the mCg and cannula placement were verified via mCherry fluorescence, and only mice with confirmed expression in the mCg→PrL pathway were included in behavioral analyses.

#### Brain slice electrophysiology of mCg→PrL projections

At P35, mice were stereotactically injected with 400 nL of rgAAV-hSyn-EGFP (1.93 × 10^13^ genomic copies/mL; #50465, Addgene), rgAAV-CaMKIIa-mCherry (2.2 × 10^13^ genomic copies/mL; #114469, Addgene), or rgAAV-CaMKIIa-EGFP (3.8 × 10¹² genomic copies/mL; #50469, Addgene) into the PrL (coordinates from bregma: AP +2.34 mm, ML ±0.4 mm, DV −1.95 mm) at a rate of 40 nL/min. Three weeks were allowed for viral expression. At P56, mice were anesthetized with isoflurane and decapitated. Brains were quickly removed, and 300-µm coronal slices of the medial prefrontal cortex were prepared using a Leica VT1200S vibratome in ice-cold, oxygenated slicing buffer (pH 7.3, 305 mOsm) containing 206 mM sucrose, 25 mM sodium bicarbonate, 2.5 mM potassium chloride, 10 mM magnesium sulfate, 1.45 mM sodium dihydrogen phosphate, 0.5 mM calcium chloride, and 11 mM D-glucose. The buffer was continuously oxygenated with 5% CO₂ and 95% O₂. Slices were transferred to a holding chamber containing oxygenated aCSF (126 mM NaCl, 26 mM NaHCO₃, 2.5 mM KCl, 1.45 mM NaH₂PO₄, 1 mM MgCl₂, 2 mM CaCl₂, 9 mM D-glucose) at room temperature (22-25 °C) for at least one hour. For recording, slices were placed in a submersion-type chamber on a microscope stage and perfused with oxygenated aCSF at 30 ± 1.5 °C using an inline heater. Recording pipettes were pulled from borosilicate glass (4-7 MΩ). Whole-cell patch-clamp recordings were performed under differential interference contrast and fluorescence microscopy targeting EGFP-positive neurons in layer V of the PrL. Spontaneous postsynaptic currents were recorded in voltage-clamp mode using SutterPatch, held at −70 mV, with compensation for electrode and cell capacitance. Recordings started five minutes after achieving whole-cell configuration and lasted five minutes. Only neurons with stable access resistance (<20% change) and baseline currents were included. Synaptic events were detected and analyzed with SutterPatch software; mean amplitude and frequency for each neuron were used for statistics. For spontaneous excitatory postsynaptic current experiments, 20 μM bicuculline was added to the bath. Intracellular solution contained 117 mM K-gluconate, 13 mM KCl, 10 mM HEPES, 2 mM Na₂ATP, 0.4 mM Na₂GTP, 1 mM MgCl₂, 0.1 mM EGTA, 0.07 mM CaCl₂, pH 7.3, 290 mOsm. Action potential firing was recorded using current injection from −50 to +490 pA in 20 pA steps; rheobase was the minimal current eliciting action potentials. For spontaneous inhibitory postsynaptic current experiments, 20 μM DNQX and 50 μM D-APV were added to isolate GABAergic currents. Intracellular solution contained 140 mM CsCl, 10 mM HEPES, 2 mM Na₂ATP, 0.4 mM Na₂GTP, 1 mM MgCl₂, 0.05 mM EGTA, 3.6 mM NaCl, pH 7.3, 290 mOsm.

#### Blood and brain collection

Blood collection and brain perfusion were performed with minor modifications to previously described methods (26, 82). To minimize the effects of circadian rhythms and acute stress from behavioral testing, blood was collected at the same time of day, 24 hours after the last behavioral session. One day following the light-dark box test, mice were anesthetized with isoflurane. One mL of blood was collected from the right atrium between 8:00 am and 11:00 am and transferred to 1.5 mL low-binding tubes. After blood collection, mice were perfused intracardially with phosphate-buffered saline (PBS) followed by 4% paraformaldehyde. Brains were extracted and post-fixed overnight in 4% paraformaldehyde at 4 °C and then transferred to a 15% sucrose solution in PBS for one day, followed by 30% sucrose for an additional day. Brains were subsequently embedded in optimal cutting temperature compound and stored at −80 °C until sectioning.

#### Immunohistochemistry

Free-floating coronal brain sections (30 μm thickness) were prepared using a Leica CM3050S cryostat and collected in PBS. Sections were blocked and permeabilized in PBS containing 5% normal goat serum (NGS) and 0.1% Triton X-100 for 1 hour at room temperature to reduce nonspecific binding. Sections were then incubated overnight at room temperature with one of the following primary antibodies: Mouse anti-CaMKIIα (1:500, Millipore, 05-532) and rabbit anti-GABA (1:1000, Sigma, A2052-25UL). After three 10-minute washes in PBS, sections were incubated with the appropriate Alexa Fluor 647-conjugated secondary antibodies (1:400, Thermo Fisher) for 2 hours at room temperature in the dark. Nuclear staining was performed with DAPI (1:50,000, Roche, 10236276001) for 3 minutes at room temperature. Sections were mounted on glass slides using Dako® fluorescent mounting medium and stored at 4 °C protected from light until imaging. Images were acquired with either a Zeiss LSM800 confocal microscope using Zen software or an Olympus BX61 epifluorescence microscope. The number of EGFP^+^, EGFP^+^CaMKIIα^+^, or EGFP^+^GABA^+^ cells was counted using Fiji/ImageJ software package (83).

#### Serum Corticosterone Measurement

Blood samples were allowed to clot at room temperature for 1 hour and then centrifuged at 3,000 rpm for 15 minutes at 4 °C. The resulting serum was carefully collected and transferred into 1.5 mL low-binding microcentrifuge tubes, then stored at –80 °C until analysis. Serum corticosterone concentrations were measured using a commercially available ELISA kit (Cayman Chemical, #501320) according to the manufacturer’s instructions with minor modifications (26, 82).

### Statistical Analysis

All statistical analyses were performed using SPSS 28 (IBM) and GraphPad Prism 8. Experiments were randomized whenever possible. For continuous data, assumptions of normality and homogeneity of variances were assessed using the Shapiro-Wilk test and Levene’s test, respectively. For experiments involving multiple groups or repeated measures, one-way analysis of variance (ANOVA), two-way ANOVA, or mixed-design ANOVA (including at least one paired measure) was performed, followed by relevant pairwise comparisons with Bonferroni correction. Data that violated only the assumption of homogeneity of variances were analyzed using Welch’s t-test or Welch ANOVA. For data that violated assumptions of normality and/or homogeneity, non-parametric tests were applied, including the Mann-Whitney U test or Kruskal-Wallis test. Statistical significance was defined as p < 0.05. Comparisons between two groups were conducted using either Student’s t-test or paired Student’s t-test, as appropriate. Effect sizes are reported for all analyses, and additional statistical parameters are provided in **Supplementary Table 1**. Data are presented as mean ± standard error of the mean (SEM).

## Acknowledgments

This work was supported by the National Institutes of Health (NIH) R01 MH-116869 (M.N.), the UAB Comprehensive Neuroscience Center Pilot Award (M.N. and S.-i.K.), the Hill Crest Foundation (M.N.), JST PRESTO grant JPMJPR14M6 (M.N.), and UAB Department of Psychiatry startup funds (M.N.). R.T. is supported by a scholarship from the Uehara Memorial Foundation. We thank members of the Niwa and Kano laboratories for their technical assistance, data analysis support, and insightful discussions. We also thank Dr. Akira Sawa and members of his laboratory for helpful discussions during the initial phase of the project. Figures 3A, 3D, 4A, 6A, and 6D were created with BioRender.com.

## Supporting Information for

**Supplemental Figure 1.**
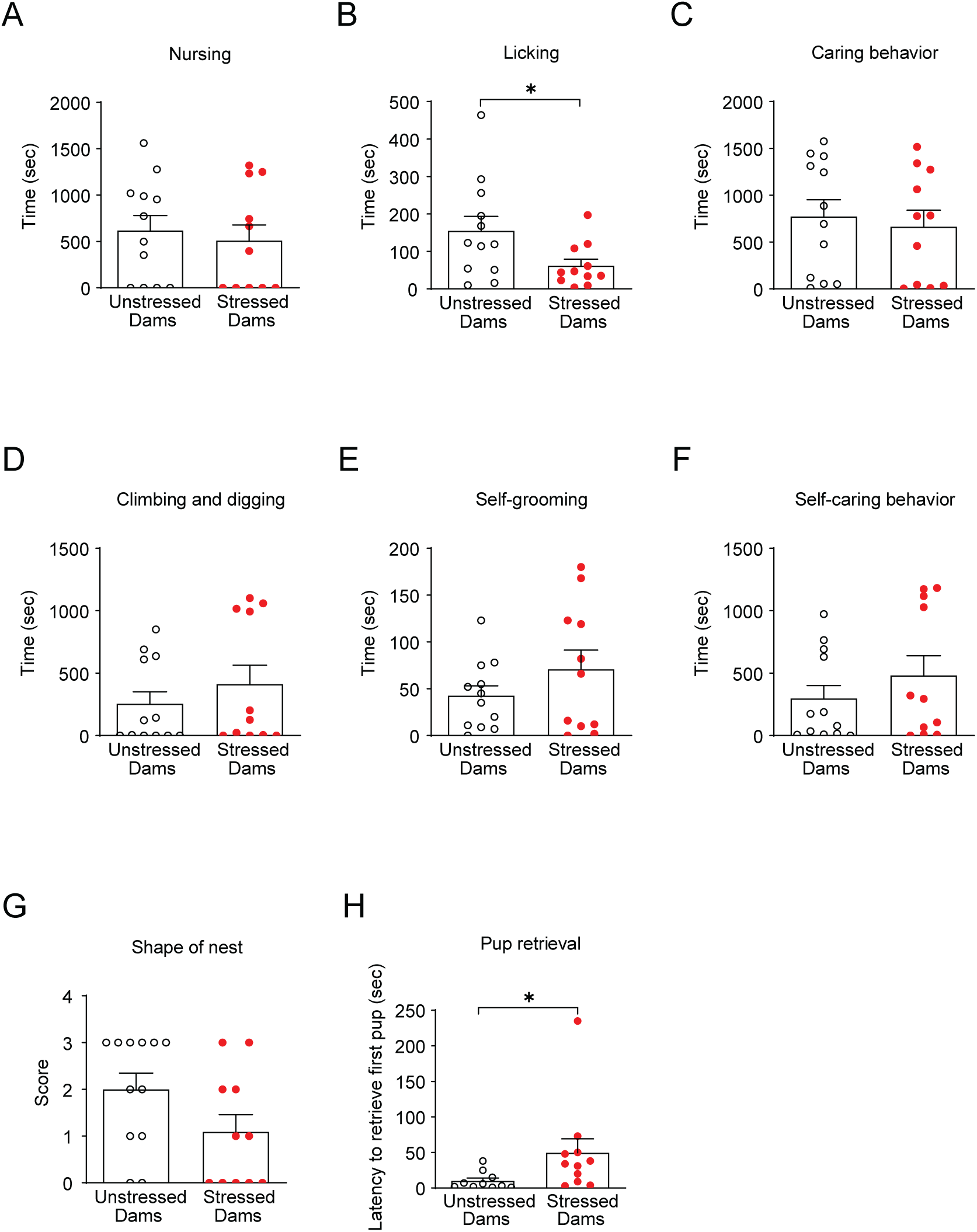
Early postpartum maternal care deficits are limited at delivery. (A-C) Caring behavior toward pups shortly after delivery (postpartum day 1) was largely unchanged, except for pup licking. (D-F) Self-care behaviors, including climbing/digging and self-grooming, were not affected. (G) Nest scores were similar between unstressed and stressed dams. (H) Pup retrieval latency was increased in stressed dams, indicating reduced maternal responsiveness. Data are shown as means ± SEM, N = 11-12 per group. *p < 0.05. Statistical tests: Mann-Whitney U (A-D, F-H) and Welch’s t-tests (E) were applied as appropriate. Detailed statistics are provided in **Supplementary Table 1**.

**Supplemental Figure 2.**
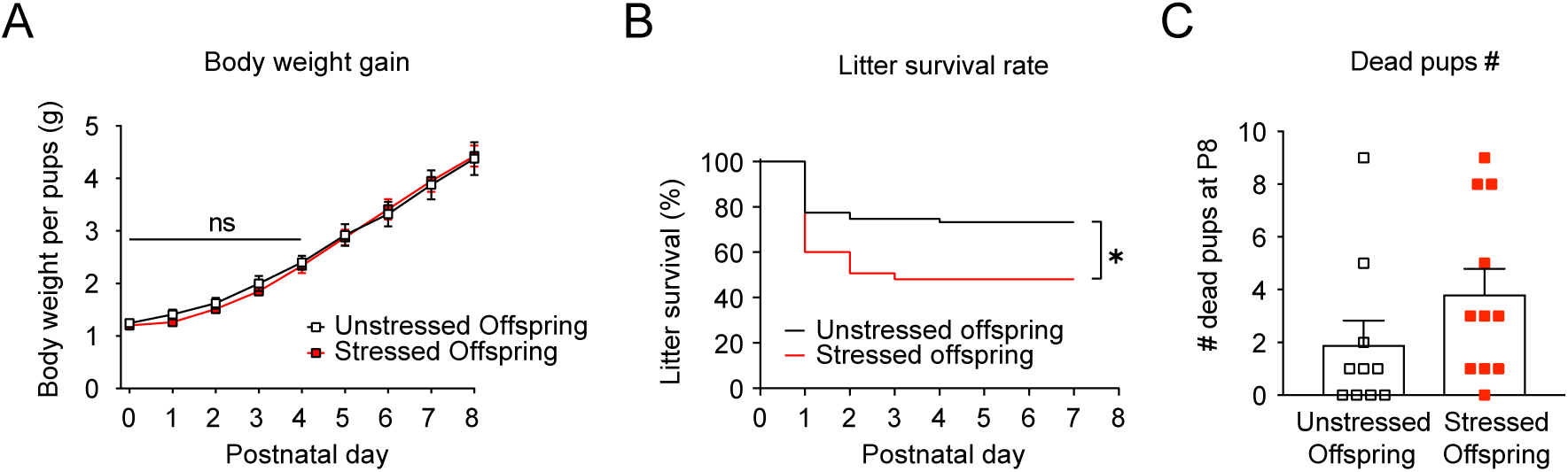
Effects of impaired maternal caregiving on offspring growth and survival. (A) Pup body weight gain over the first postnatal week was similar between groups. (B) Survival rates were reduced in stressed offspring compared to unstressed controls. (C) Number of dead pups per litter was not significantly different, although stressed litters tended to have more deaths. Data are shown as means ± SEM, N = 10-11 litters per group. *p < 0.05. Statistical tests: Mann-Whitney U and Student’s t Test (A, C) and Log-rank Mantel-Cox (B) were applied as appropriate. Detailed statistics are provided in **Supplementary Table 1**.

**Supplemental Figure 3.**
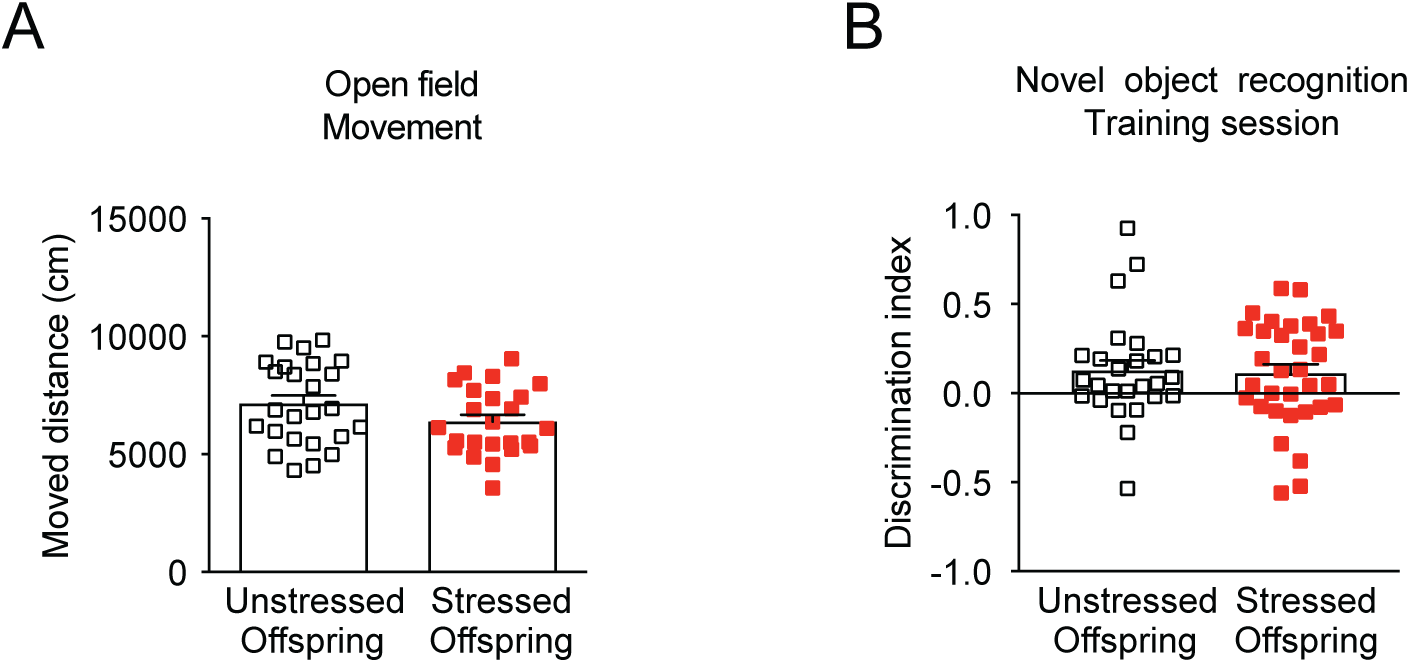
Validation of behavioral measures. (A) Total distance moved in the open field test was similar between groups, indicating no differences in locomotor activity. (B) Discrimination index in the novel object recognition test during the training session was similar across groups. Data are shown as means ± SEM, N = 24-25 per group. *p < 0.05. Statistical tests: Student’s t-test (A) and Mann-Whitney U test (B) were applied as appropriate. Detailed statistics are provided in **Supplementary Table 1**.

**Supplemental Figure 4.**
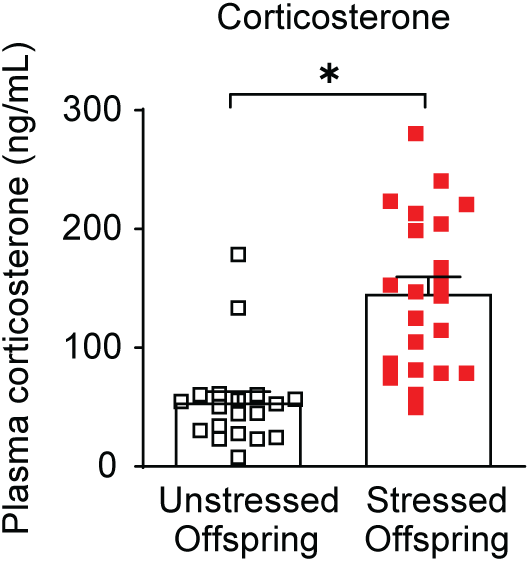
Increased plasma corticosterone levels in stressed offspring. Plasma corticosterone levels were elevated in adult offspring reared by stressed dams. Data are shown as means ± SEM, N = 19-22 per group. *p < 0.05. Statistical tests: Mann-Whitney U test was applied. Detailed statistics are provided in **Supplementary Table 1**.

**Supplemental Figure 5.**
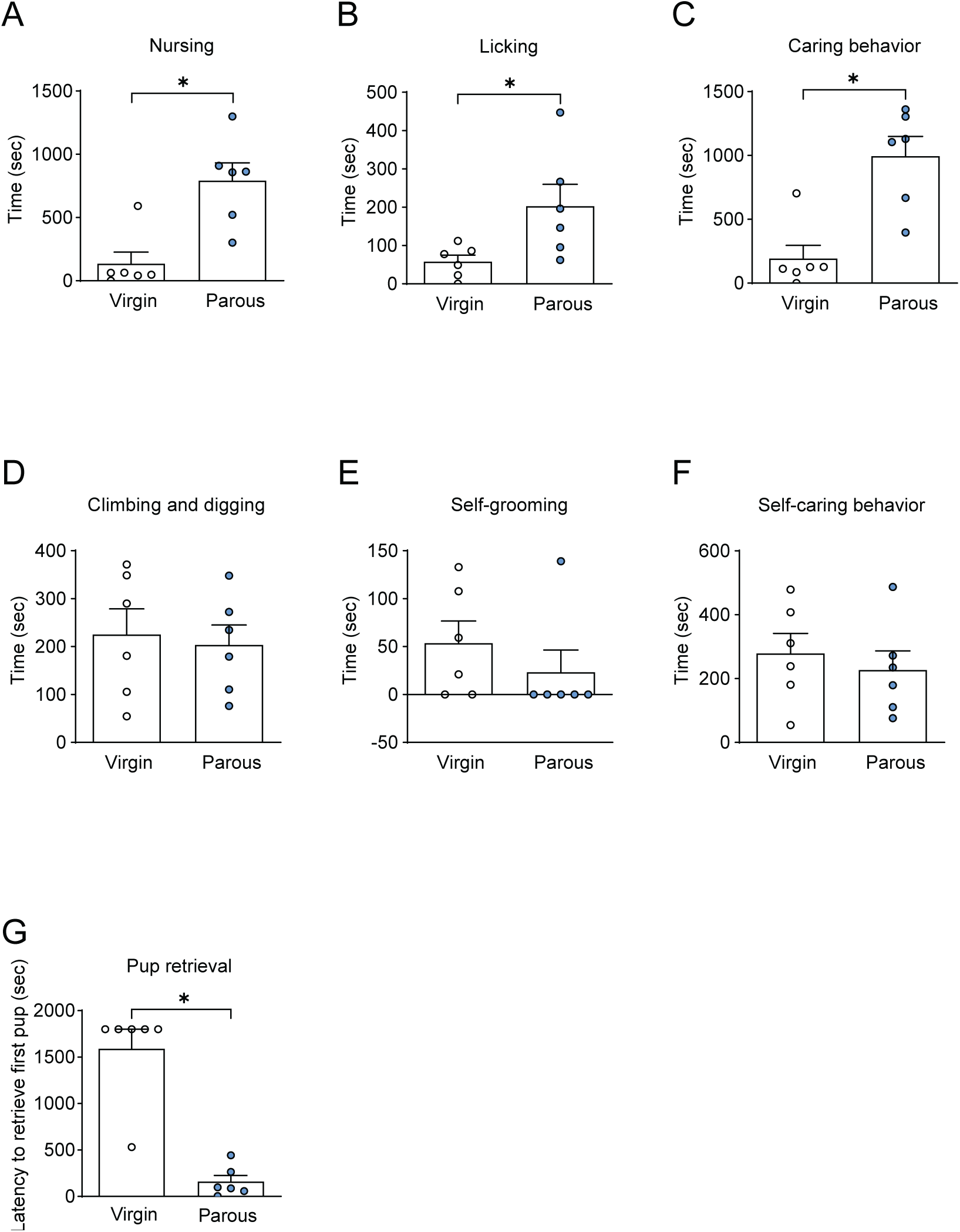
Parous mice display robust maternal behavior. (A-C) Pup-directed behaviors, including nursing, licking, and retrieval, were increased in parous females compared to virgins. (D-F) Self-care behaviors were similar between virgin and parous females. (G) Pup retrieval latency was reduced in parous females, reflecting enhanced maternal responsiveness. Data are shown as means ± SEM, N = 6 per group. *p < 0.05. Statistical tests: Mann-Whitney U test (A, C, E, G) and Student’s t-test (B, D, F) were applied as appropriate. Detailed statistics are provided in **Supplementary Table 1**.

**Supplemental Figure 6.**
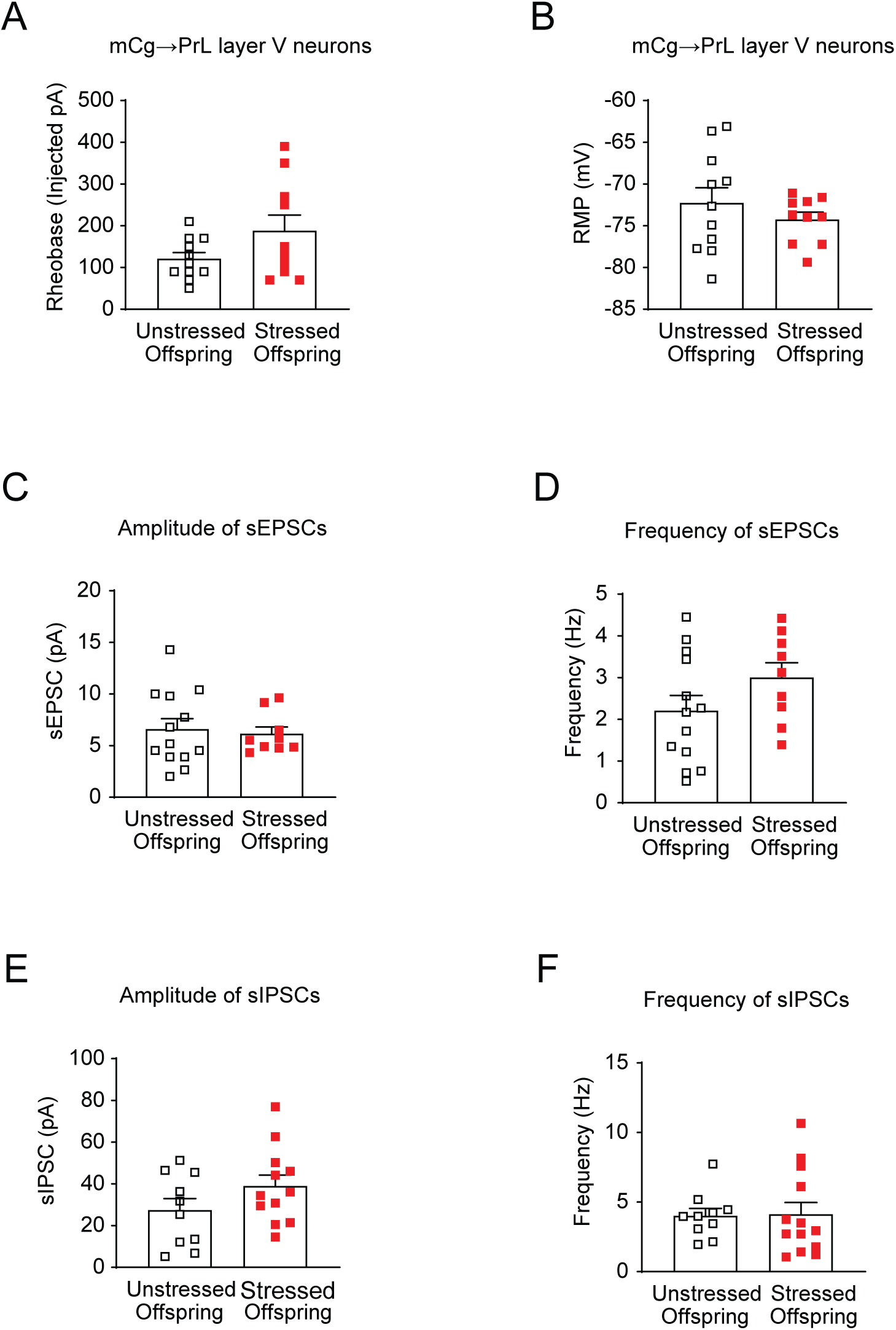
Electrophysiological characterization of mCg→PrL neurons in stressed offspring. (A) Rheobase of these neurons showed no change. (B) Resting membrane potential was similar between groups. (C-F) Amplitude and frequency of spontaneous excitatory and inhibitory currents were unchanged, indicating intact presynaptic input. Data are shown as means ± SEM, N = 10-12 cells per group. Each group included cells obtained from at least three animals. Statistical tests: Two-way mixed ANOVA (A) and Student’s/Welch’s t-tests (B-G) were applied. Detailed statistics are provided in **Supplementary Table 1**.

**Supplementary Table 1.**
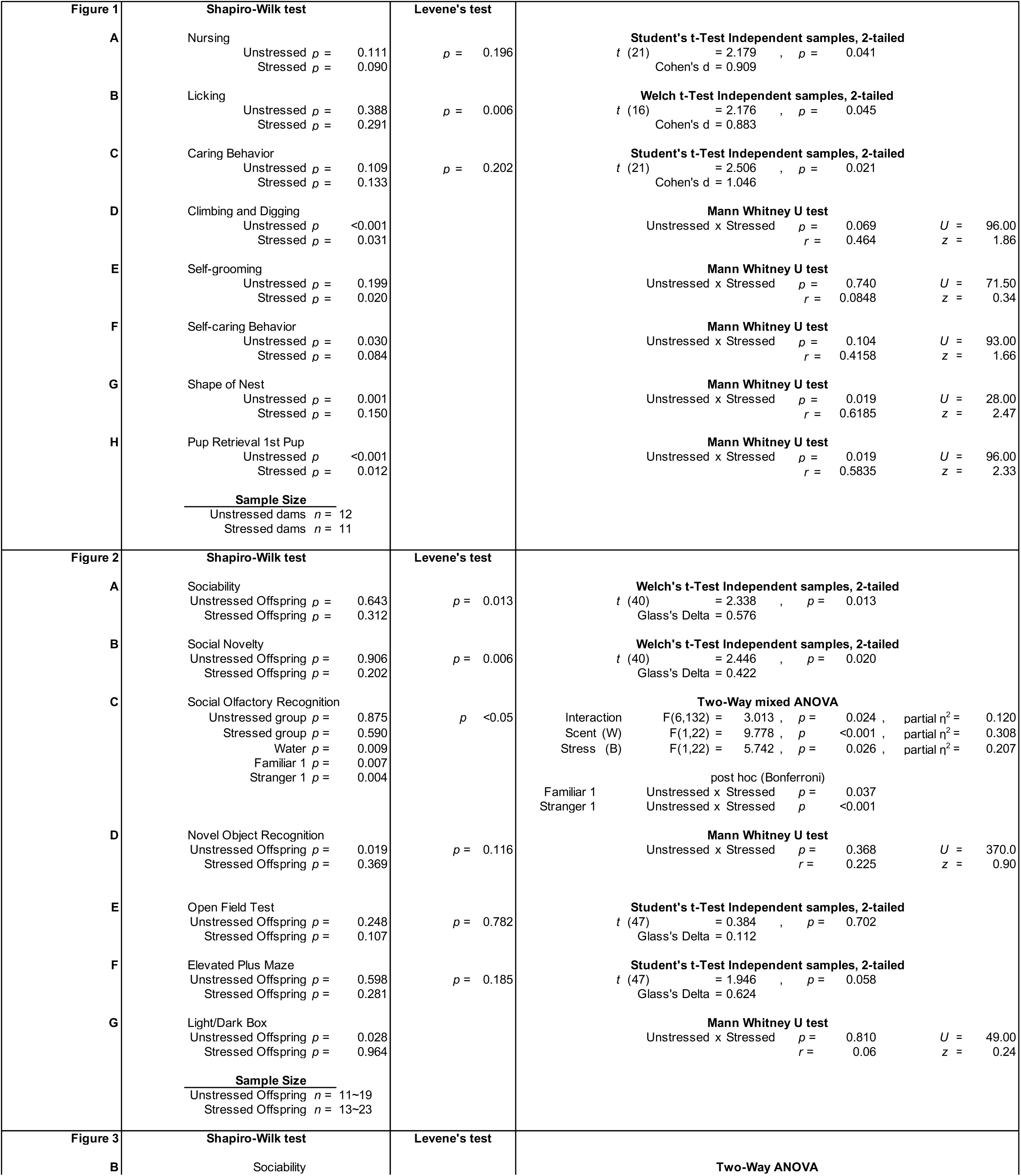
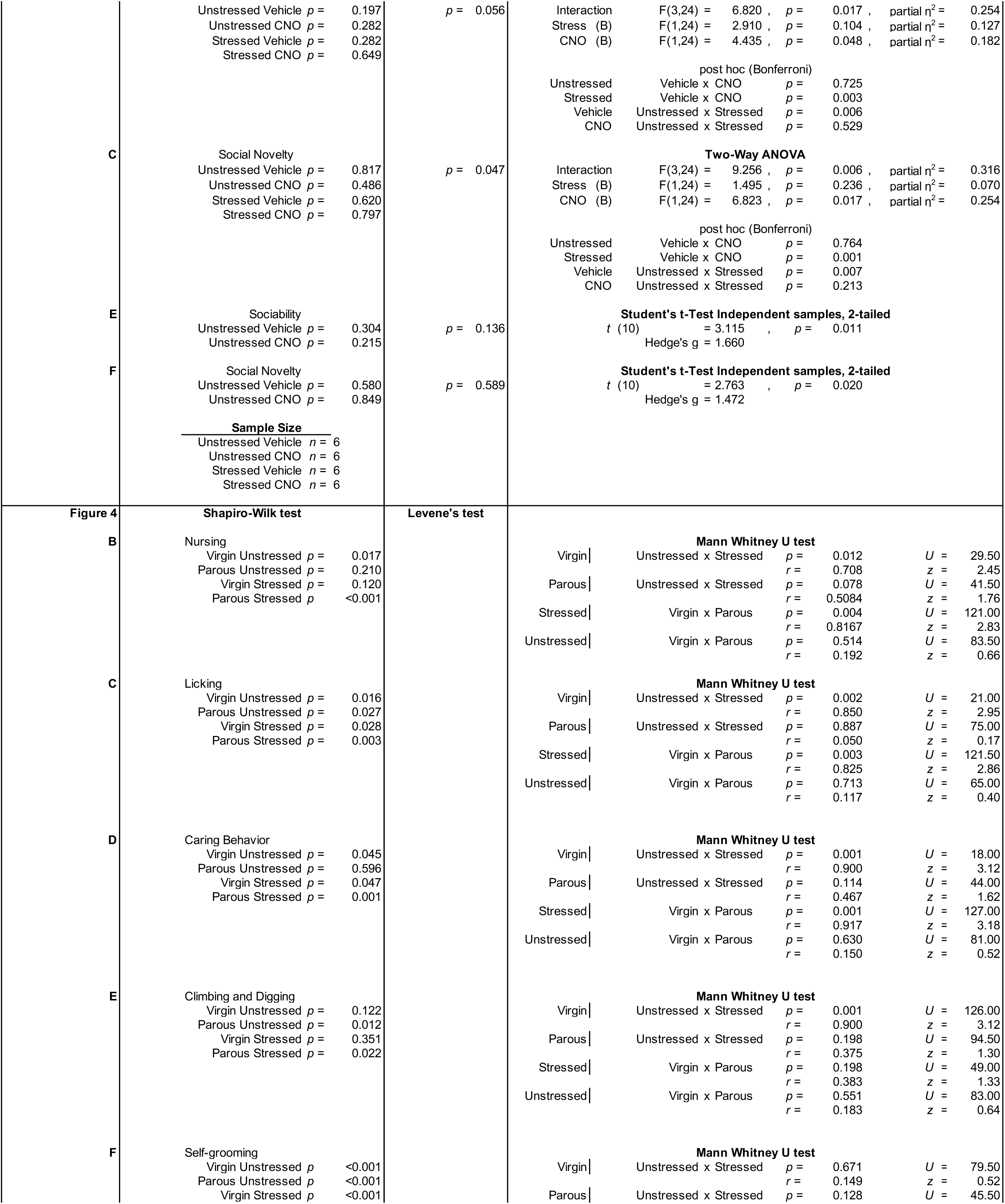
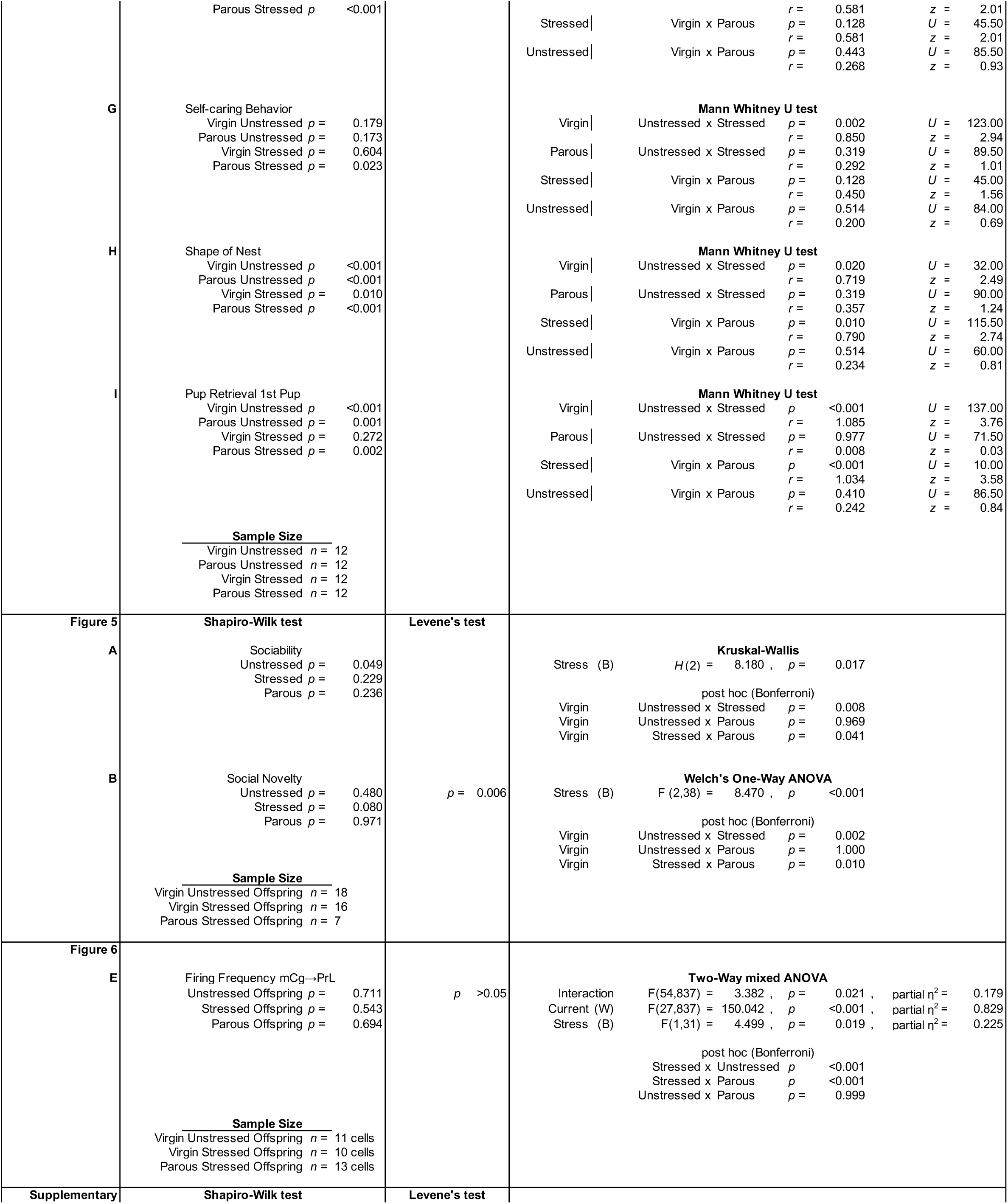
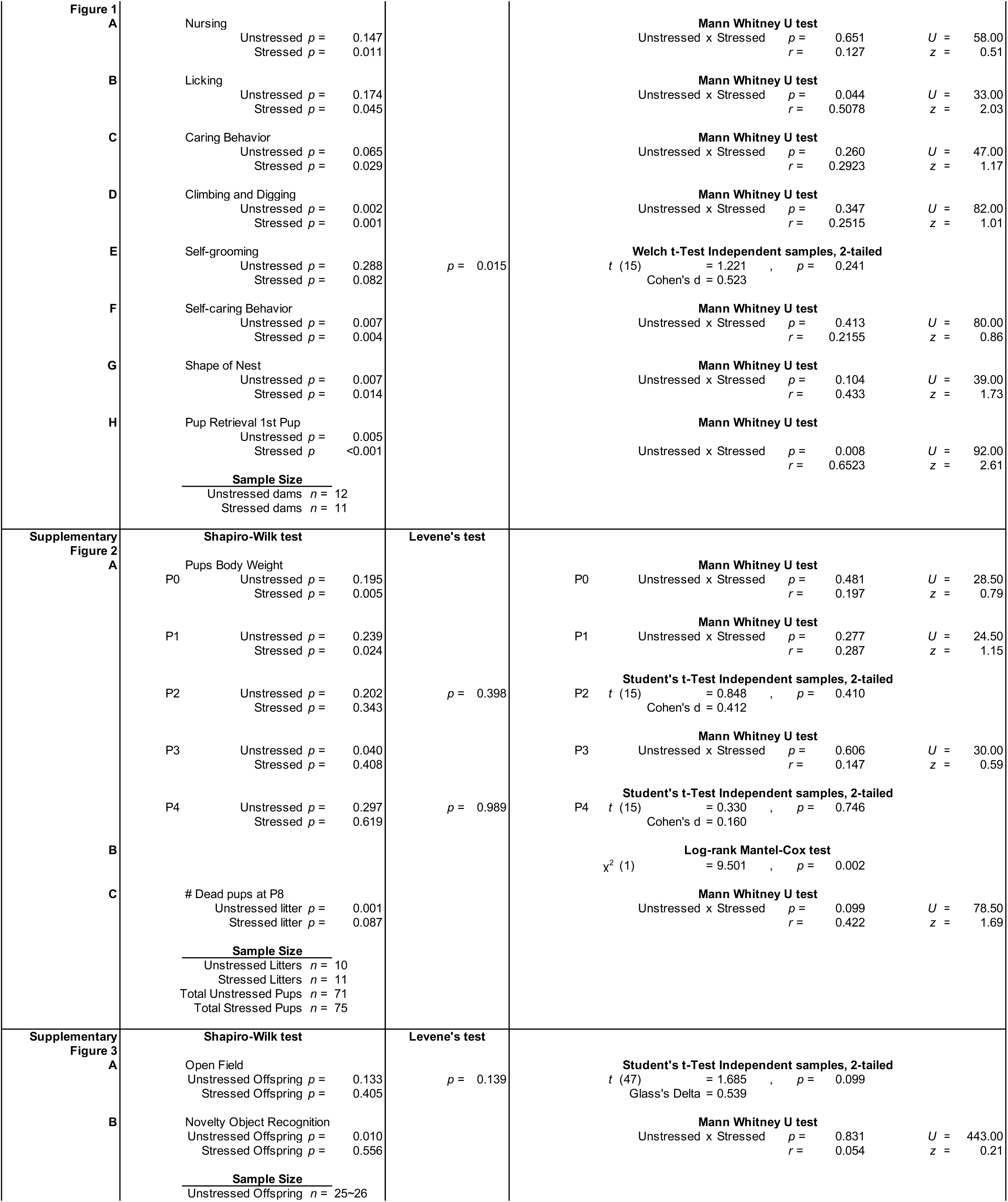
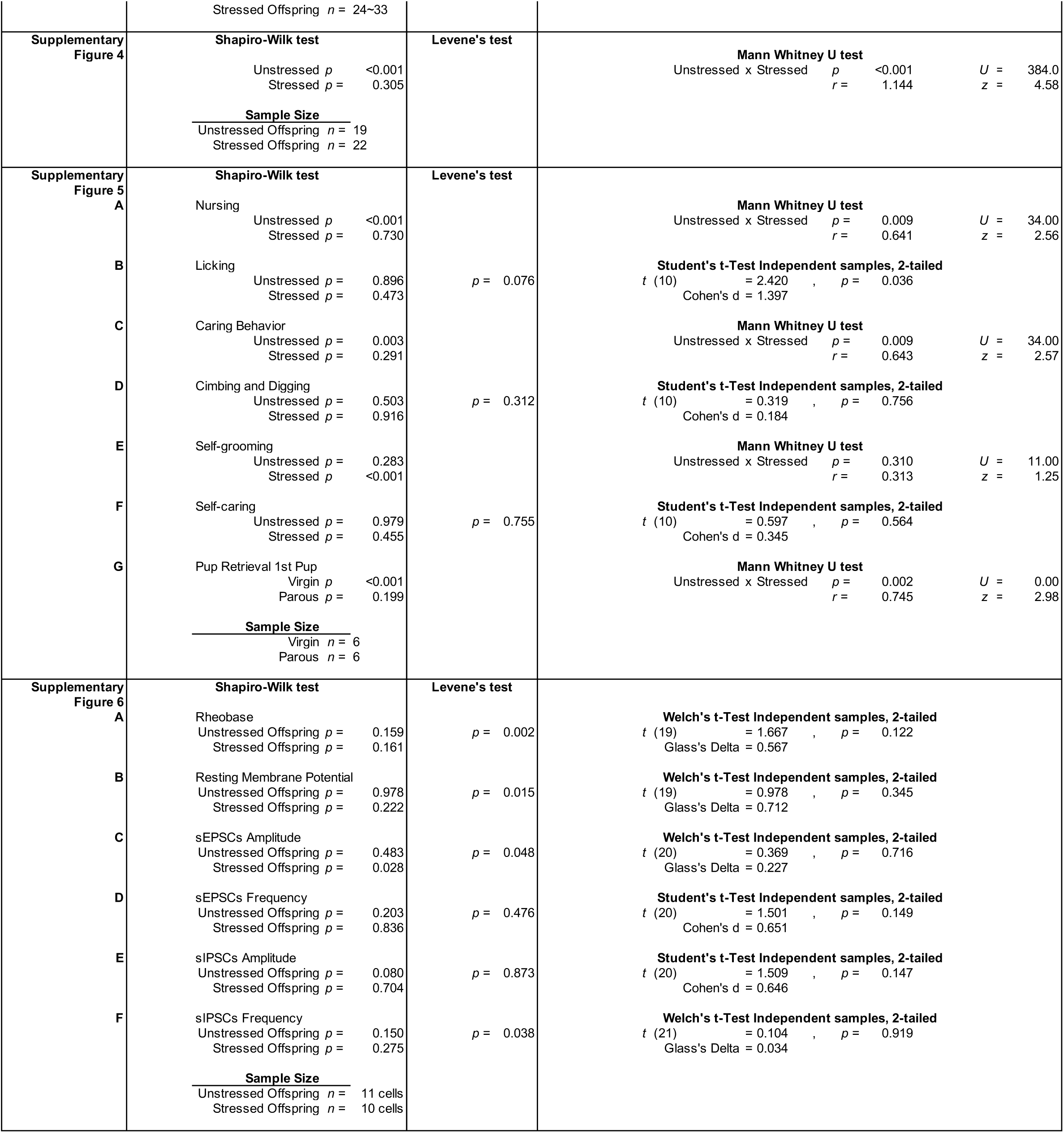
Statistical analyses for all results. For each dataset, sample sizes, normality tests, statistical tests performed, *p* values, effect sizes, and post hoc comparisons are reported. For repeated-measures ANOVAs, selected within-group factors were included in post hoc analyses.

